# Redundancy Circuits of the Commissural Pathways in Human and Rhesus Macaque Brains

**DOI:** 10.1101/2020.09.03.281931

**Authors:** Zulfar Ghulam-Jelani, Jessica Barrios-Martinez, Aldo Eguiluz-Melendez, Ricardo Gomez, Yury Anania, Fang-Cheng Yeh

## Abstract

It has been hypothesized that the human brain has traded redundancy for efficiency, but the structural existence has not been identified to examine this claim. Here, we report three redundancy circuits of the commissural pathways in primate brains, namely the orbitofrontal, temporal, and occipital redundancy circuits of the anterior commissure and corpus callosum. Each redundancy circuit has two distinctly separated routes connecting a common pair of cortical regions. We mapped their trajectories in human and rhesus macaque brains using individual and population-averaged tractography. The dissection results confirmed the existence of these redundancy circuits connecting the orbitofrontal lobe, amygdala, and visual cortex. The volume analysis showed a significant reduction in the orbitofrontal and occipital redundancy circuits of the human brain, whereas the temporal redundancy circuit had a substantial organizational difference between the human and rhesus macaque. Our overall findings suggest that the human brain is more efficient in the commissural pathway, as shown by the significantly reduced volume of the anterior commissure which serves as the backup connections for the corpus callosum. This reduction of the redundancy circuit may explain why humans are more vulnerable to psychiatric brain disorders stemming from the corpus callosum compared to non-human primates.

**Significance:** We report and describe the connection routes of three redundancy circuits of the commissural pathways in human and rhesus macaque brains and compare their volumes. Our tractography and dissection studies confirmed that the human brain has smaller redundancy circuits. This is the first time such redundancy circuits of the commissural pathways have been identified, and their differences quantified in human and rhesus macaque to verify the redundancy-efficiency tradeoff hypothesis. The findings provide new insight into the topological organization of the human brain and may help understand the circuit mechanism of brain disorders involving these pathways.

## Introduction

It has been hypothesized that the human brain has traded-off redundancy for efficiency to allocate connectivity for higher cognitive functioning such as intelligence (Marsman et al, 2017). Recently, Pryluk et al. highlighted that human neurons have more efficient encoding than monkeys (Pryluk et al., 2019). They suggested that humans have traded off robustness for efficiency and hypothesized that the human brain may be more susceptible to psychiatric disorders. At the macroscopic level, the efficiency of a brain network can be defined and quantified using the graph analysis (Bullmore and Sporns, 2009). A recent structural connectome study using a population-averaged template of young adults has shown that commissural pathways—the anterior commissure and corpus callosum—contribute to most of the brain network efficiency due to their long-range connections between the two hemispheres (Yeh et al., 2018). Studies have also shown that functional connections are organized as a highly efficient system in the human brain (Sporns et al, 2004; Stam, 2004; Eguiliuz et al., 2005; Achard et al., 2006; van den Heuvel et al., 2008). These findings suggest a relationship between efficiency and intelligence. However, there is no structural connectome evidence showing that the topological organization of the human brain is more efficient or has traded-off robustness with efficiency.

To investigate this efficiency trade-off hypothesis at the commissural pathways, here we focus on a specific structural organization called the “redundancy circuit.” We define a redundancy circuit as a network of the brain structure that includes two distinctly separated white matter pathways connecting two common functional areas to form a fail-safe connection to ensure robustness (Fig. 1a). The separated white matter connections may increase neuronal synchronization, as was observed in Pryluk’s study for monkeys (Pryluk et al., 2019). If one of the connections has an injury or agenesis, information exchange may not be interrupted entirely as an alternative pathway may be present (Fig. 1b). This redundancy connection thus has greater robustness against dysconnectivity. In this study, we hypothesize that primate brains may have structural existence of a redundancy circuit of the commissural pathways and that the human brain may have a smaller redundancy connection that trades off structural robustness for efficiency.

**Figure 1:**
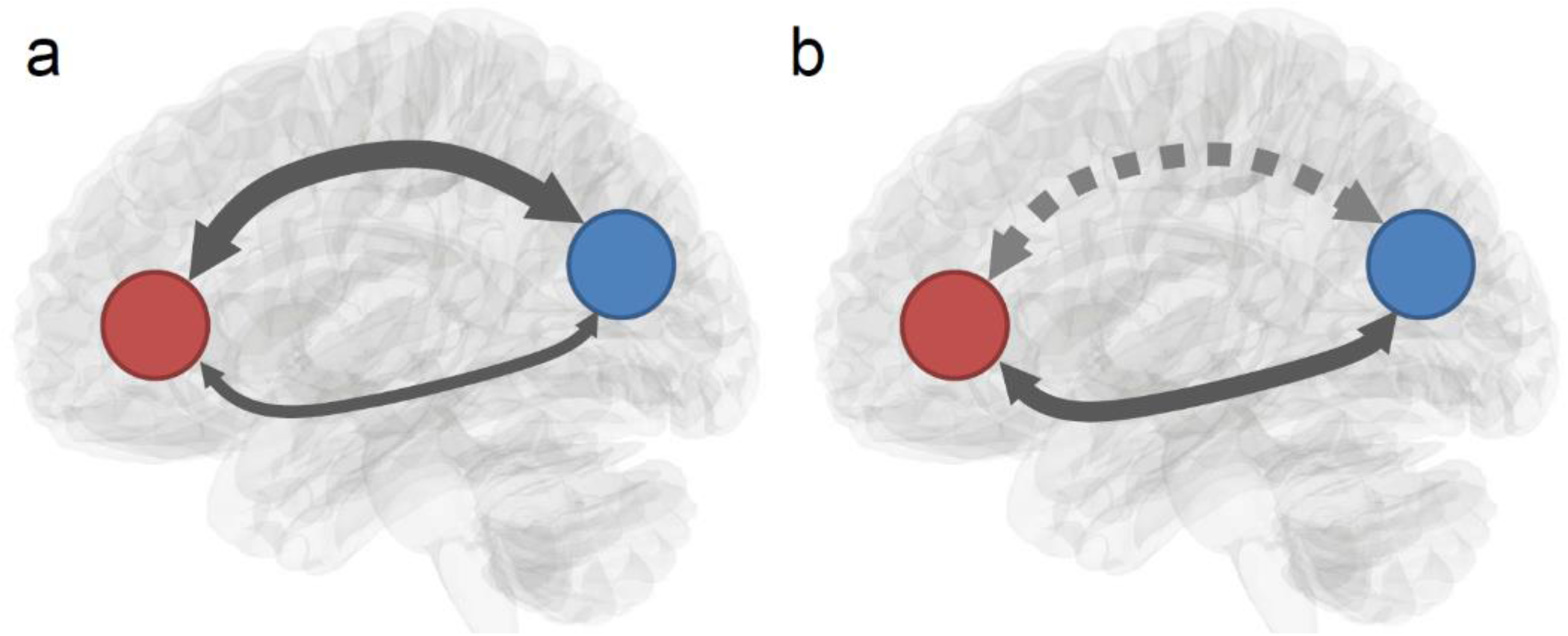
Schematic illustration of a redundancy circuit. (a) A redundancy circuit has two distinctly separated white matter pathways connecting two common functional areas to form robustness in connectivity. (b) An injury or agenesis of one white matter pathway in the redundancy circuit may disrupt one connection. However, the information exchange between the innervated regions is not entirely interrupted due to the existence of an alternate pathway connecting to them.

To examine these hypotheses, we performed diffusion magnetic resonance imaging (MRI) fiber tracking in *in-vivo* human and rhesus macaque brains as well as in a population-averaged template of 1021 young human adults and a population-averaged template of 36 rhesus macaques. We limited our search to the anterior commissure and corpus callosum to identify the existence of redundancy circuits due to our previous study showing that the commissural pathways contribute to most of the network efficiency in the human brain (Yeh et al., 2018). Trajectories of the identified redundancy circuits were then validated using cadaveric dissection to provide the tissue evidence. We then compared the volumetric differences in the redundancy circuits between the human and rhesus macaque to examine the efficiency trade-off hypothesis and investigated whether the redundancy circuits of the commissural pathways in the human brain are significantly reduced in comparison with those of the rhesus macaque.

## Results

### Redundancy circuits in the human brain

Our study based in population-averaged tractography identified three redundancy circuits of the commissural pathways in the human brain (Figure 2). Figure 2a shows the axial view of three redundancy circuits we identified in the population-averaged tractography of 1021 young human adults. The three redundancy circuits are named orbitofrontal, temporal, and occipital redundancy circuits, respectively. All of them have two distinctly separate connection routes contributed from the corpus callosum and the anterior commissure, matching our definition of a redundancy circuit. Tractography results show that the orbitofrontal redundancy circuit includes the orbitofrontal branches of the anterior commissure (red) and the forceps minor of the corpus callosum (dark red). Both branches connect to the orbitofrontal cortex. The temporal redundancy circuit includes the temporal branch of the anterior commissure (green) and the tapetum of the corpus callosum (dark green). Both branches connect to the amygdala and medial temporal lobe. The occipital redundancy circuit includes the occipital branch of the anterior commissure (blue) and the forceps major of the corpus callosum (dark blue). Both connect to the visual cortex in the occipital lobe. The redundancy circuits in the human brain is also visualized in the supplementary materials (Fig. S2).

**Figure 2:**
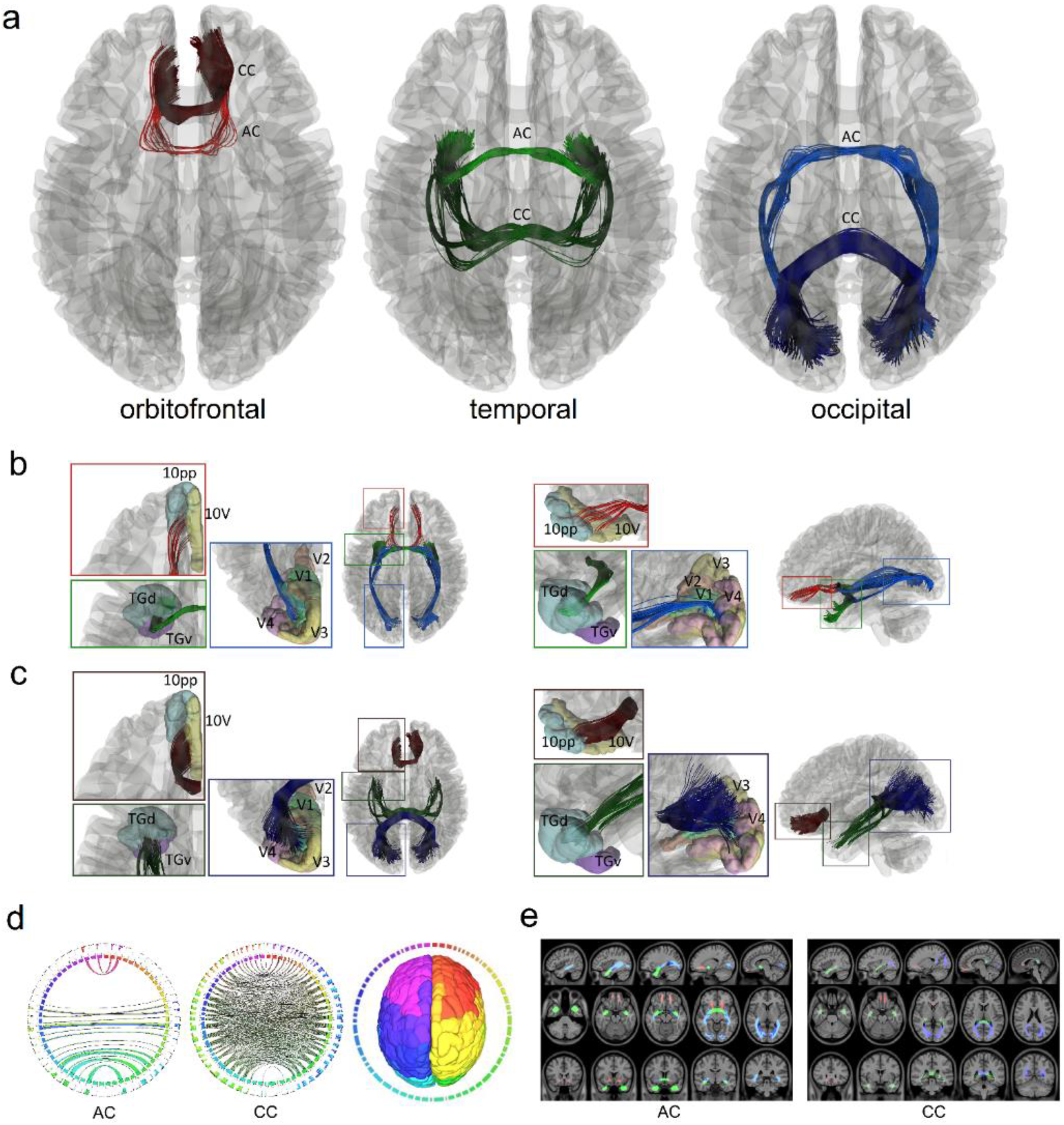
Redundancy circuits of the commissural pathways identified in the human brain. (a) The tractogram shows the fiber tracts composition of the orbitofrontal (red), temporal (green), and occipital (blue) redundancy circuits. Each of them connects two cortical regions with two distinctly different connection routes through anterior commissure and the corpus callosum. (b) The two tractograms show the anterior commissure components of the three redundancy circuits in axial and sagittal views. The orbitofrontal component (red) connects between left-right 10pp and 10v. The temporal (green) component connects primarily to the left and right TGd and TGv region. The occipital (blue) component connects V1 primarily, and partly to V2, V3, and V4. (c) The two tractograms show the corpus callosum components of the three redundancy circuits in axial and sagittal views. The three components (red, green, blue) also connect to the same cortical regions of their anterior commissure counterpart but through a completely different route in the midsagittal plane. (d) Connectograms represents bilateral connectivity patterns of the average human anterior commissure and corpus callosum, confirming that these two commissure pathways connect to common parcellated region pairs. (e) The white matter map of the anterior commissure and corpus callosum forming the redundancy circuits confirms that their connections travel through entirely different routes. AC = anterior commissure, CC = corpus callosum.

We further zoom in at the cortical regions connected by these redundancy circuits. The cortical segmentation was defined using the HCP-MMP atlas (Glasser et al., 2016). The abbreviation of the connecting regions is listed in the supplementary materials (Table S1). Figure 2b shows the anterior commissure portion of the redundancy circuits. The red-colored tract is part of the orbitofrontal redundancy circuit that projects anteriorly to the orbitofrontal cortex and connects primarily to the polar 10p region (10pp) and area 10v region (10v). The green-colored tract is part of the temporal redundancy circuit that projects laterally to the amygdala and connects primarily to the dorsal temporal polar cortex subregion (TGd). The tract also projects near the ventral temporal polar cortex subregion (TGv). The blue-colored tract is part of the occipital redundancy circuit that connects primarily to the primary visual cortex (V1) and partly projects to the secondary, tertiary, and quaternary visual cortices (V2, V3, and V4). Figure 2c shows the corpus callosum portion of the three redundancy circuits. The dark red-colored tract corresponds to the branch of the corpus callosum that project to the orbitofrontal cortex, the dark green-colored tract to the amygdaloidal cortex, and the dark blue-colored tract to the visual cortex. Similar to the pathways in the anterior commissure, the corpus callosal branches connect to the same cortical regions, including 10pp and 10v (orbitofrontal redundancy circuit), TGd and TGv (temporal redundancy circuit), and V1, V2, V3, and V4 (occipital redundancy circuit). Although connecting regions are the same, the connection trajectories are projected from different white matter routes.

We further confirmed the existence of redundancy circuits of the commissural pathways using connectograms (Fig. 2d) for both the anterior commissure and corpus callosum. The connectograms also confirmed that both the anterior commissure and portions of the corpus callosum share the same connecting targets between the left and right hemispheres. Figure 2e shows the voxelwise location of the redundancy circuits in axial, sagittal, and coronal views for the anterior commissure and corpus callosum. The images show that target regions are connected by entirely separated routes. This fulfills our definition of a redundancy circuit that two cortical regions are connected by two separate white matter tracts to form connection redundancy (Fig. 1).

### Redundancy circuits in the rhesus macaque brain

We also identified three similar redundancy circuits of the commissural pathways in the rhesus macaque brain. Figure 3 shows these redundancy circuits mapped by population-averaged tractography. Fig. 3a shows the axial view of three redundancy circuits in the rhesus macaque brain of a population template averaged from 36 rhesus macaque brains. The overall fiber topology is the same as for the human shown in Fig. 2. The orbitofrontal redundancy circuit includes the orbitofrontal branches of the anterior commissure (red) and the forceps minor of the corpus callosum (dark red). The temporal redundancy circuit includes the temporal branch of the anterior commissure (green) and the tapetum of the corpus callosum (dark green). Both branches connect to the amygdala and medial temporal lobe. The occipital redundancy circuit includes the occipital branch of the anterior commissure (blue) and the forceps major of the corpus callosum (dark blue), both connecting to the visual cortex in the occipital lobe. The redundancy circuits of the human and rhesus macaque show similar fiber topology. The redundancy circuits of the commissural pathways in the rhesus macaque brain is also visualized in the supplementary materials (Fig. S3).

**Figure 3:**
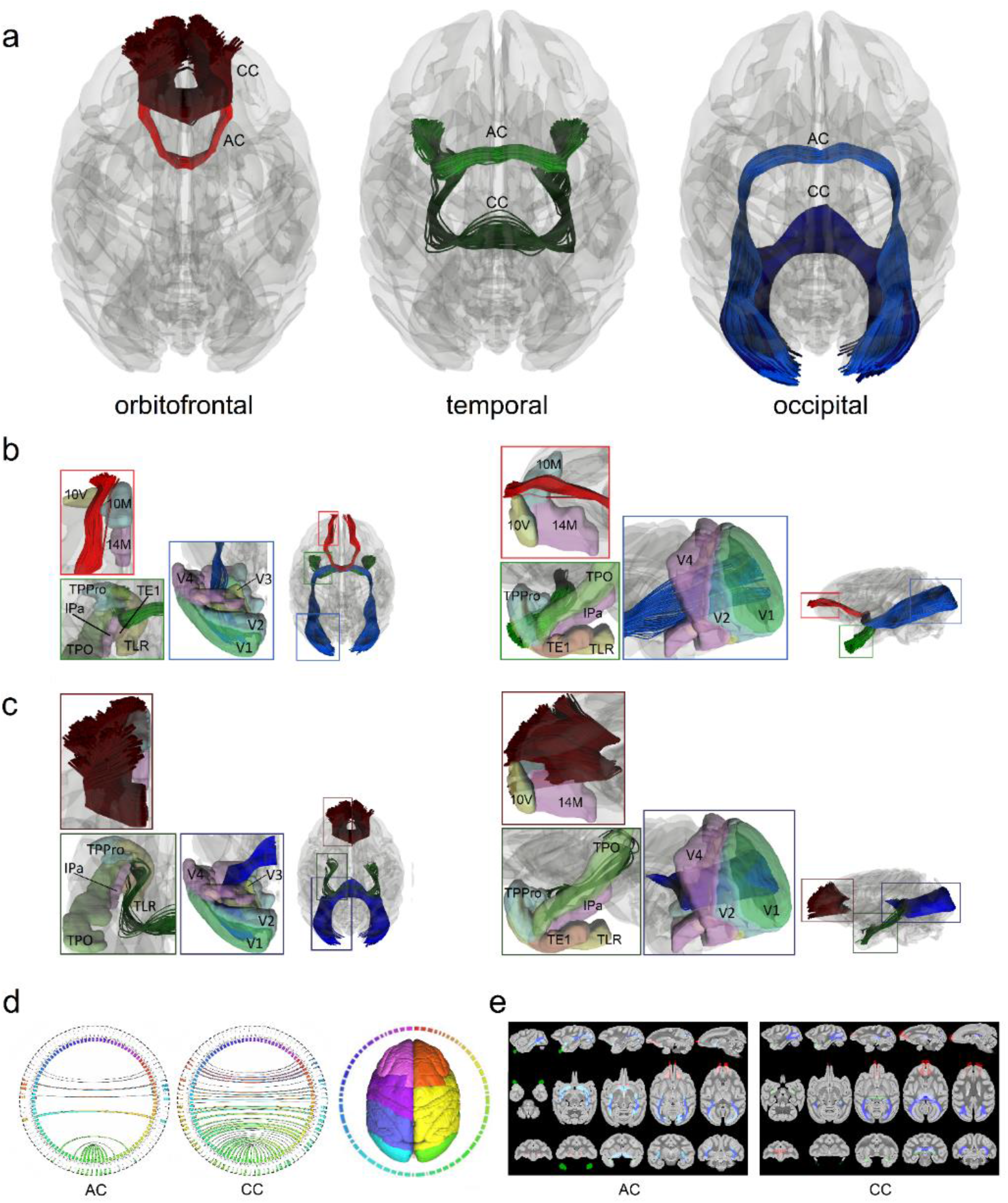
Redundancy circuits of the commissural pathways identified in the rhesus macaque brain. (a) The tractogram shows the fiber tracts composition of the orbitofrontal (red), temporal (green), and occipital (blue) redundancy circuits. Each of them connects two cortical regions with two distinctly different connections route through anterior commissure and the corpus callosum. (b) The two tractograms show the anterior commissure components of the three redundancy circuits in axial and sagittal views. The orbitofrontal component (red) connects between left-right 10M and 10V, while some connections are to 14M. The temporal (green) component connects primarily to the left and right TPPro region while other fibers continue to TLR(R36) and TPO regions. The occipital (blue) component connects V1 primarily. (c) The two tractograms show the corpus callosum components of the three redundancy circuits in axial and sagittal views. The three components (red, green, blue) also connect to the same cortical regions of their anterior commissure counterpart but through a completely different route in the midsagittal plane. (d) Connectograms represents bilateral connectivity patterns of the average human anterior commissure and corpus callosum, confirming that these two commissure pathways connect to common parcellated region pairs. (e) The white matter map of the anterior commissure and corpus callosum forming the redundancy circuits confirms that their connections travel through entirely different routes. AC = anterior commissure, CC = corpus callosum.

More detailed connecting regions of the left hemisphere in the sagittal and axial views are illustrated in Fig. 3b and Fig. 3c for the anterior commissure and corpus callosum, respectively. The cortical segmentation is defined using the CIVM rhesus atlas (Calabrese et al., 2015). The abbreviation of the connecting regions is listed in the supplementary materials (Table S1). Figure 3b shows the anterior commissure portions of three redundancy circuits of the commissural pathways. The red-colored tract is a part of the orbitofrontal redundancy circuit that projects anteriorly to the orbitofrontal cortex and connects primarily to the medial and ventral parts (10M and 10V) of area 10 cortex while some connections are made within area 14 of the medial part (14M). The green-colored tract is part of the temporal redundancy circuit that projects laterally to the amygdala and connects primarily to the temporopolar prisocortex (TPPro) region while other fibers continue to the area TL rostral part (TLR(R36)) and temporal parieto-occipital associated area in TPO. The blue-colored tract is part of the occipital redundancy circuit and connects primarily to the primary visual cortex (V1) and partly projects to the secondary, tertiary, and quaternary visual cortices (V2, V3, V4). Figure 3c shows the corpus callosum portion of the three redundancy circuits of the commissural pathways. The dark red-colored tract corresponds to the branch of the corpus callosum that project to the orbitofrontal cortex, the dark green-colored tract to the amygdaloidal cortex, and the dark blue-colored tract to the visual cortex. Similar to the pathways in the anterior commissure, the corpus callosum branches connect to the same cortical regions including the 10M, 10V, and 14M (orbitofrontal redundancy circuit), TPPro, TLR(R36), and TPO (temporal redundancy circuit), and V1, V2, V3, and V4 (occipital redundancy circuit). We further confirm the existence of redundancy circuits using connectograms (Fig. 3d) and the voxelwise projection (Fig. 3e) of the pathways. Similar to the results in the human brain, the connectograms confirmed that both the anterior commissure and part of the corpus callosum share the same connecting targets between the left and right hemispheres. The voxelwise projection images of the pathways label the white matter regions occupied by the redundancy circuits in axial, sagittal, and coronal views. The images show that the target regions are connected by entirely separated routes. Again, this fulfills our definition of a redundancy circuit that a common brain region is connected by two separated white matter tracts to form connection redundancy as we described for the human brain.

### Cadaveric dissections of the anterior commissure

We validated our tractography findings using cadaveric dissections. Here, we did not include the corpus callosum as neuroanatomy literature has covered its structure substantially (Mooshagian, 2008; Goldstein et al., 2019). Figure 4 shows the dissection results of the anterior commissure for the human brain, and Figure 5 shows the dissection results of the anterior commissure for the rhesus macaque brain. The dissections were completed by a neuroanatomist (Dr. Aldo Eguiluz-Melendez) and labeled by three different neuroanatomists (Dr. Aldo Eguiluz-Melendez, Dr. Ricardo Gomez, and Dr. Yury Anania). The different branch projections of the anterior commissure are color-coded with dotted lines to show the orbitofrontal branch (FB, red), temporal branch (TB, green), and the occipital branch (OB, blue). The abbreviations are listed in the supplementary materials (Table S2). The inset images with the dissection pictures show the orientation of the brain. Overall, the fibers in Fig. 4 and 5 show a topology that is content with our tractography findings (Fig. 2 and Fig. 3). In the human brain, the red dotted lines in Fig. 4 show the orbitofrontal branch (FB) of the anterior commissure branching off from the main trunk in the anterior/inferior thalamus (Fig. 4a, 4c, and 4d) and projecting anteriorly to the lower frontal lobe (Fig. 4a). The remaining bundles contain the temporal branch (TB, green) and occipital branch (OB, blue), as shown in Fig. 4d and 4f. These two branches separate in the anterior temporal lobe (Fig. 4a, 4b, 4d, and 4f). The temporal branch (TB) bends anteriorly toward the amygdaloidal region (Fig. 4a, 4d, and 4f), whereas the occipital branch (OB) turns posteriorly (Fig. 4a and 4d) and projects toward the visual cortex (Fig. 4a and 4e). Similarly, in the rhesus macaque brain, the red dotted lines in Fig. 5 show the orbitofrontal branch (FB) of the anterior commissure branching off from the main trunk in the anterior/inferior thalamus (Fig. 5a, 5b, 5c, and 5d) and projecting anteriorly to the lower frontal lobe (Fig. 5a and 5b). The separation location of the temporal and occipital branch in the human and the rhesus macaque are the same. The separation of the temporal branch (TB, green) and the occipital branch (OB, blue) occurs in the anterior temporal lobe (Fig. 5a, 5b, 5d, 5e, and 5f). The temporal branch (TB) bends anteriorly toward the amygdaloidal region (Fig. 5a, 5b, and 5d), while the occipital branch (OB) turns posteriorly (Fig. 5a, 5b, and 5f) and projects towards the visual cortex (Fig. 5a, 5b, 5e, and 5f). The dissection results provided tissue evidence confirming the trajectories and termination location of the three redundancy circuits identified by tractography. Moreover, the dissection results also show that the rhesus macaque brain has a relatively larger volume of the occipital redundancy circuits compared to the human brain.

**Figure 4:**
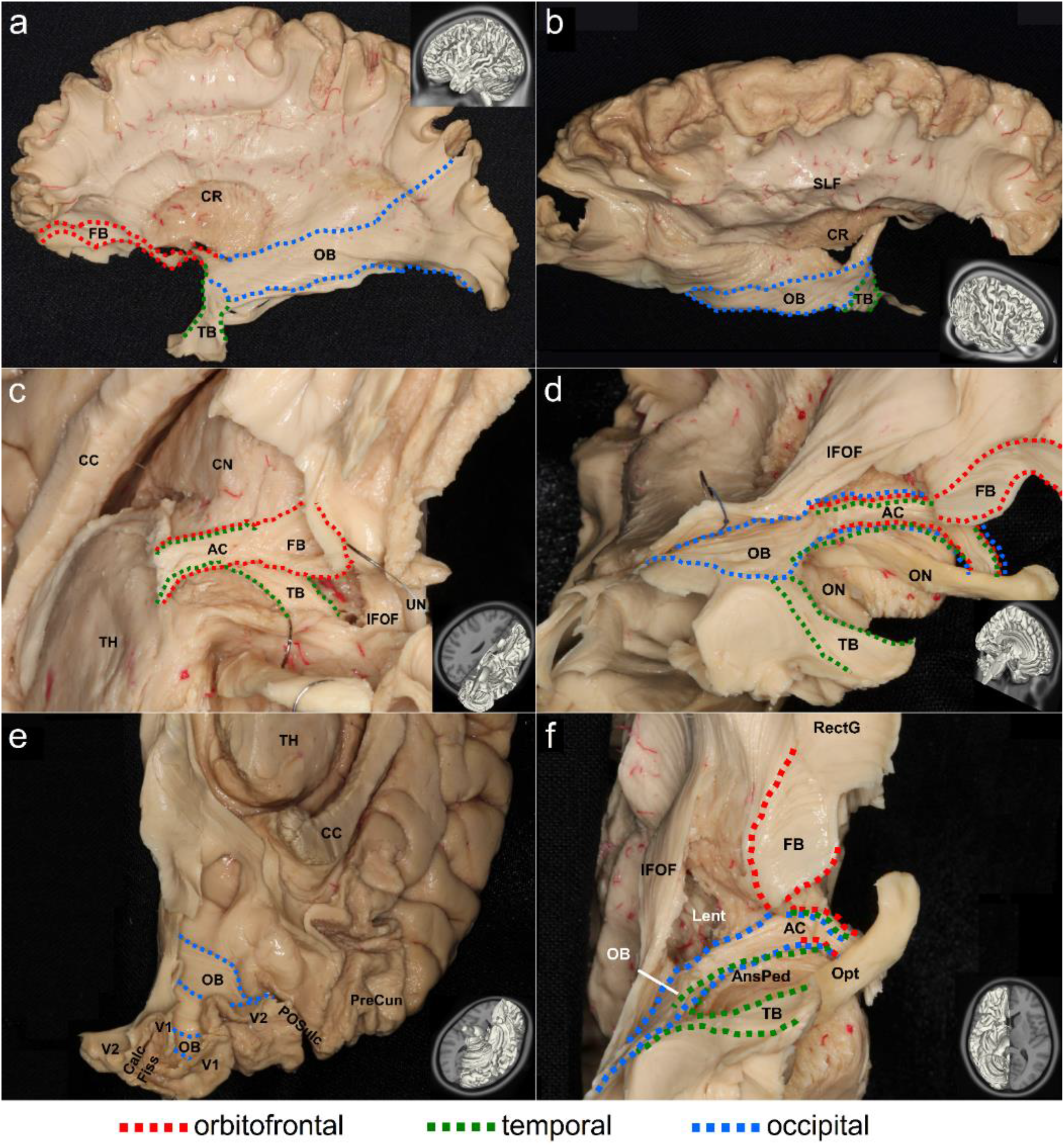
Cadaveric dissection of the anterior commissure that is part of the redundancy circuits of the commissural pathways in the human brain. The different branches of the anterior commissure are shown in dotted lines with red indicating projections to the orbitofrontal regions, green indicating connections to the temporal regions, and blue indicating projections to the occipital regions. The dissection photo views the anterior commissure from left-lateral (a), right-anterior-lateral (b), and inferior-lateral views (c,d,e,f) of the anterior commissure. The orbitofrontal branch travels through the midsagittal plane and connects the bilateral lower orbitofrontal regions (c,d,f). The temporal branch also travels through the same midsagittal region and connects the bilateral amygdaloid region (c,d,f). The occipital branch similarly travels through the midline and connects the bilateral visual cortices (d,e,f). The connecting route of the fiber pathways for each redundancy circuit matches the findings from diffusion magnetic resonance imaging (MRI) fiber tracking.

**Figure 5:**
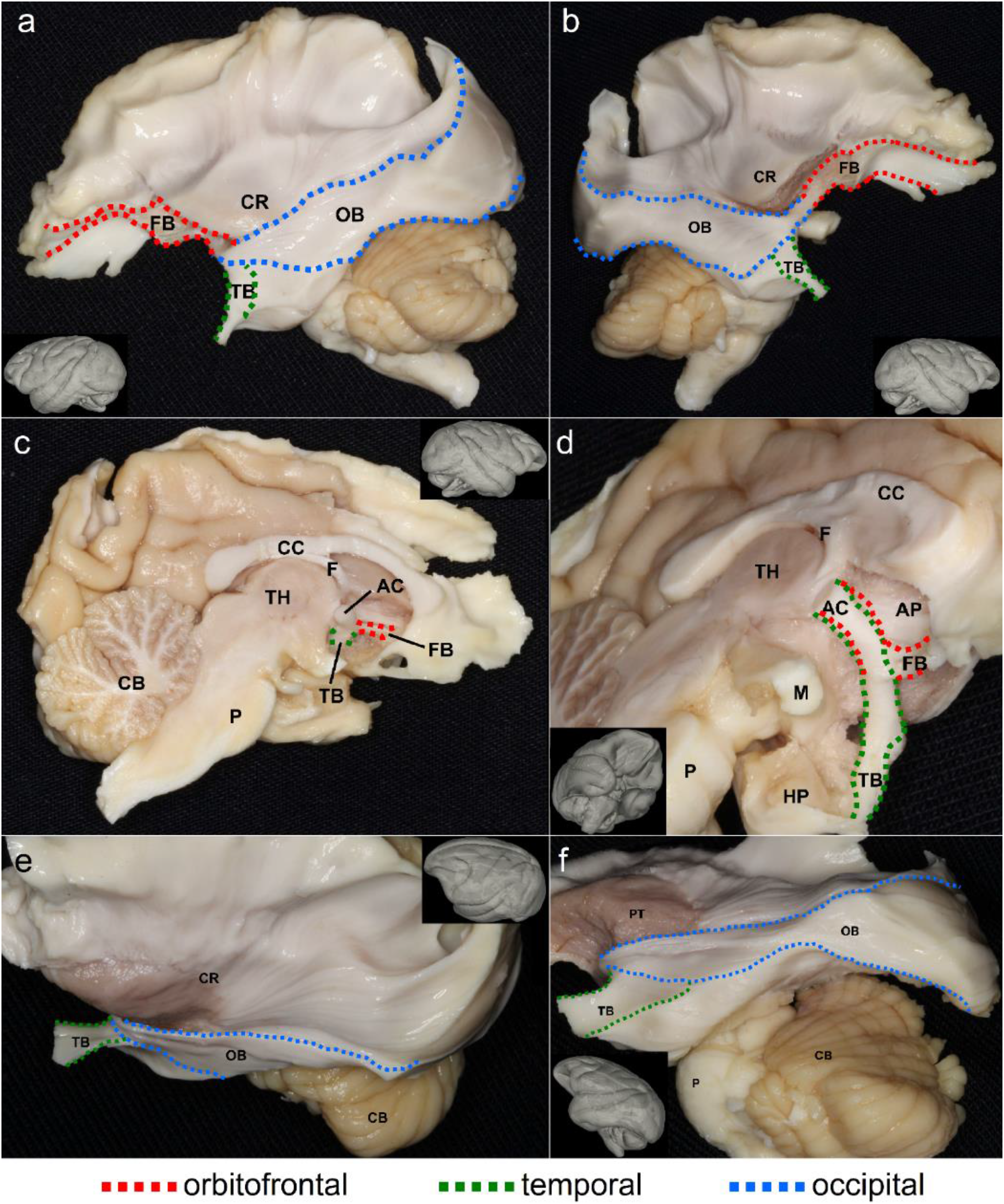
Cadaveric dissection of the anterior commissure that is part of the redundancy circuits of the commissural pathways in the rhesus macaque brain. The different branches of the anterior commissure are shown in dotted lines with red indicating projections to the orbitofrontal regions, green indicating connections to the temporal regions, and blue indicating projections to the occipital regions. The dissection photo views the anterior commissure from left-lateral (a, e, f), right-lateral (b), right-midsagittal (c), and inferior-lateral (d) views of the anterior commissure. The orbitofrontal branch travels through the midsaggital plane and connects the bilateral lower orbitofrontal regions (a,b,c,d). The temporal branch also travels through the same midsagittal region and connects the bilateral amygdaloid region (a,b,c,d). The occipital branch similarly travels through the midline and connects the bilateral visual cortices (a,b,e,f). The connecting route of the fiber pathways for each redundancy circuit matches the findings from diffusion magnetic resonance imaging (MRI) fiber tracking.

### Volumetric comparison between the human and rhesus macaques

We further performed a volumetric comparison between the human and rhesus macaque brains to test our hypothesis that the human brain has less redundancy in the connection circuit of the commissural pathways. As illustrated in Fig. 6, by using the tract volume fraction quantified by the total volume of tract divided by brain volume. We obtained the volume fraction using 27 human subjects and 30 rhesus macaque subjects (Materials and Methods). As shown in Fig. 6a, the volume fraction of the anterior commissure component (AC), the corpus callosum component (CC), and their sum (AC+CC) in the orbitofrontal redundancy circuit in humans were all significantly smaller than that of the rhesus macaque (p = 0.020, 0.017, and 0.013). Although the differences were statistically significant, the effect size was not large (Cohen’s d = 0.66, 0.62, and 0.69). Overall, the rhesus macaque brain had a slightly larger volume fraction of the orbitofrontal redundancy circuit in the corpus callosum than that of the human. Fig. 6b shows the volume fraction of the temporal redundancy circuits separated into the anterior commissure component (AC), the corpus callosum component (CC), and their sum (AC+CC). Only the anterior commissure component is significantly larger in the rhesus macaque than the human (p = 0.025, Cohen’s d = 0.57). Fig. 6c shows the volume fraction of the occipital redundancy circuits separated into the anterior commissure component (AC), the corpus callosum component (CC), and their sum (AC+CC). All of them were significantly and substantially smaller in the human brain (p = 1.52E-05, 5.93E-08, and 6.53E-08, Cohen’s d = 1.26, 1.55, and 1.65). Overall, all redundancy circuit of the commissural pathways presents a significant reduction in the human brain, and among these three circuits, the occipital redundancy circuit showed the most considerable reduction.

**Figure 6:**
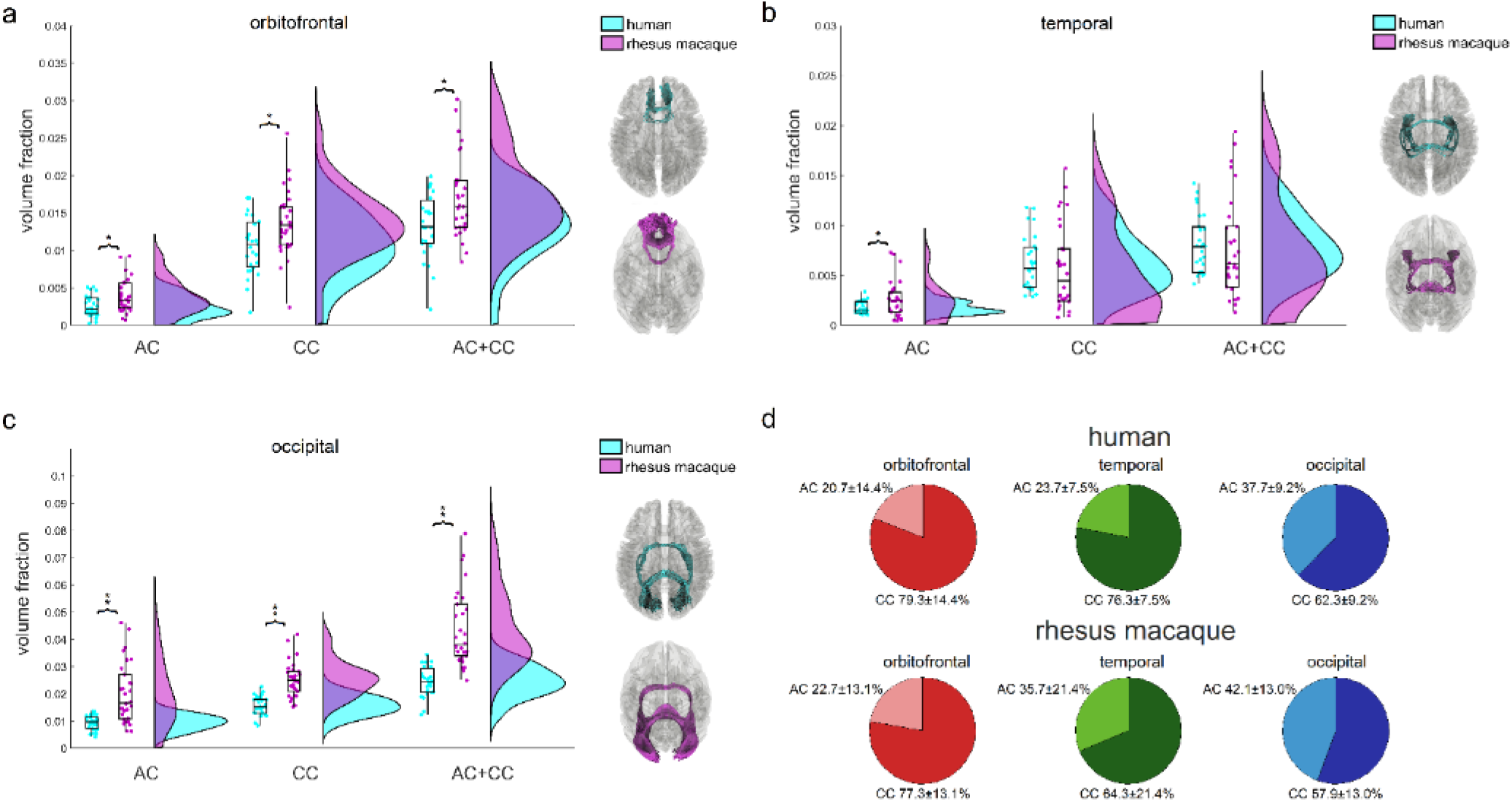
Quantitative analysis comparing the volumetrics of redundancy circuits between human and rhesus macaque commissural pathways. The volumes of the pathways are normalized against brain size to calculate the volume fraction for comparison. The box-raincloud plot shows the volume fractions for the anterior commissure component, corpus callosum component, and their sum in the orbitofrontal (a), temporal (b), and occipital (c) redundancy circuits. The human brain shows a general trend of volumetric reduction in the redundancy circuits of the commissural pathways. The orbitofrontal redundancy circuit and its two component connections are all significantly smaller in the human brain (* < 0.05), whereas the occipital redundancy circuit also shows the same findings with a greater significance (** < 0.01). In contrast, the temporal redundancy circuit only shows a smaller volume fraction of the anterior commissure component in the human (* < 0.05). (d) The pie charts show the volumetric percentages between the anterior commissure and the corpus callosum components of the redundancy circuits. The percentage in the temporal redundancy circuit shows a significant difference between human and rhesus macaque. AC = anterior commissure, CC = corpus callosum, AC+CC = combined anterior commissure and corpus callosum.

The component percentages of the redundancy circuit devoted to the anterior commissure versus the corpus callosum for the human and rhesus macaque brain are shown in Fig. 6d. Overall, the corpus callosum takes on more volume fraction in the human brain for all three redundancy circuits, particularly the temporal redundancy circuits. The orbitofrontal redundancy circuit in the human brain shows the anterior commissure and corpus callosum have component percentages of 23.7±7.5% and 76.3±7.5%, respectively, while in the rhesus macaque brain, they are 22.7±13.1% and 77.3±13.1%. The component percentage of the corpus callosum in the human brain (76.3%) is not significantly different from that of rhesus macaque (77.3%)(p = 0.597). For the temporal redundancy circuit, the component percentages are 23.7±7.5% and 76.3±7.5% for the anterior commissure and corpus callosum, respectively. For the rhesus macaque, the percentages are 35.7±21.4% and 64.3±21.4%. The component percentage of the corpus callosum in the human brain (76.3%) is significantly and substantially larger than that of rhesus macaque (64.3%)(p = 0.0074, d = 3.33). The occipital redundancy circuit has the largest contribution from the anterior commissure in comparison. The component percentages for the anterior commissure and corpus callosum are 37.7±9.2% and 62.3±9.2% in the human brain, and 42.1±13.0% and 57.9±13.0% in the rhesus macaque brain. The component percentage of the corpus callosum in the human brain (62.3%) is not significantly different from that of rhesus macaque (57.9%)(p = 0.157). In summary, the orbitofrontal and occipital redundancy circuits of the commissural pathways have no significant difference in the percentages of their component connections, whereas the temporal redundancy in the human brain has a much larger component percentage of the corpus callosum. This suggests that there could be a substantial organizational difference in the temporal redundancy circuit of the human brain.

## Discussion

Here, we report the existence of three redundancy circuits of the commissural pathways of primate brains, namely the orbitofrontal, temporal, and occipital redundancy circuits. Each circuit has two distinctly different connecting routes contributed from parts of the anterior commissure and corpus callosum. The structural existence of these redundancy circuits in primates allows us to make a comparison to examine whether human brains have organized toward efficiency or robustness.

### The human brain is more “efficient” in the commissural pathways

Our comparison of the volume ratio of the commissural pathways confirmed a significant reduction of redundancy circuits in the human brain, confirming our hypothesis that the structural topology of the human brain is organized to be more efficient in the commissural pathways. Further, component percentage analysis showed that the temporal redundancy circuits have a significant organizational difference between human and rhesus macaque, with the human brain showing a much larger component percentage in the corpus callosum. The larger percentage of the corpus callosum may also indicate a more efficient organization, as our previous study has shown the role of the corpus callosum in the efficiency of the brain network (Yeh et al., 2018), and the presence of the corpus callosum indicates a superior form of interhemispheric communication due to its shorter distance between the dorsal parts of the isocortex (Johnson et al., 1982, 1994). In this sense, the anterior commissure could be a remnant fiber tract and has mostly been replaced by the more efficient corpus callosum (Winter and Franz, 2014), and the decrease of the anterior commissure in the human would allow for the corpus callosum to improve brain efficiency in information coding.

### Anterior commissure: a backup connection

The anterior commissure and corpus callosum have been studied in patients with severed commissural pathways or agenesis of the corpus callosum, in which the corpus callosum and hippocampal commissure are naturally not present. It has been previously reported that for some cases, an enlarged anterior commissure may help compensate for the absence of the corpus callosum in individuals with callosal agenesis (Barr and Corballis, 2002). The neuronal axons may be re-routing to achieve better functional compensation. In cases of callosal agenesis, the anterior commissure is often found to be hypertrophied (Lemire et al., 1975) and may take on the role of substituting the transfer of information (Fischer et al., 1992; Barr and Corballis, 2002). This further suggests that the connections to the severed corpus callosum could be replaced with connections to the anterior commissure with the role that redundancy circuits of the commissures play for the brain. This indicates that the recruitment of the redundancy circuit can compensate for the role of the main circuit (i.e., the corpus callosum). However, the substitution cannot fully replace the role of the corpus callosum—it is not able to transfer information for spatial analysis in the case of callosal agenesis (Martin, 1985). Moreover, in certain cases, the redundancy circuits of the commissural pathways do not take over the role of the main circuit.

### Redundancy circuit and brain diseases

The reduction of the anterior commissure in humans may leave the corpus callosum more vulnerable to dysconnectivity risk and might be the reason humans more commonly present these types of psychiatric disorders than non-human primates. This argument may be supported by the close relationship between the corpus callosum and psychiatric disorders (Bhatia et al., 2016, Brambilla et al., 2004, David et al., 1993, Swayze et al., 1990). The existence of an occipital redundancy circuit may explain why schizophrenic patients have fewer visual hallucinations than audio hallucinations, as no redundancy circuits are innervating the primary auditory cortex in the commissures. Understanding the role that the commissures and the redundancy circuits play in the human brain can help us disentangle the sophisticated mechanism behind psychiatric disorders, most of which still have much unknown in the causality, and lead us to possible prevention or treatment for patients with these disorders.

## Acknowledgments

We thank Jennifer L. Collinger, Steven Chase, and Ferdows Juya for their critical review of the manuscript. The research reported in this publication was supported by NIMH under award number R56MH113634. The content is solely the responsibility of the authors and does not necessarily represent the official views of the National Institutes of Health. Data were provided in part by the Human Connectome Project, WU-Minn Consortium (Principal Investigators: David Van Essen and Kamil Ugurbil; 1U54MH091657) funded by the 16 NIH Institutes and Centers that support the NIH Blueprint for Neuroscience Research; and by the McDonnell Center for Systems Neuroscience at Washington University.

## Author Contributions

Z.G., J.B., and F.Y. performed analyses. A.E. performed cadaveric dissections. A.E., R.G., and Y.A. labeled dissection images. Z.G., J.B., and F.Y. wrote the manuscript.

## Materials and Methods

### Human MRI experiments

We conducted a subject-specific deterministic fiber tractography study in 30 right-handed, neurologically healthy male and female subjects with age range 23–35. The data came from the Human Connectome Project (HCP) online database [WU-Minn Consortium (Principal Investigators: David van Essen and Kamil Ugurbil; 1U54MH091657)] funded by the 16 NIH institutes and centers that support the NIH Blueprint for Neuroscience Research and by the McDonnell Center for Systems Neuroscience at Washington University (Van Essen et al., 2013). Likewise, data from 1021 individual HCP subjects were utilized to compile the averaged diffusion atlas.

For the human fiber tracts, a multishell diffusion scheme was used, and the b-values were 1000, 2000, and 3000 s/mm^2^. The number of diffusion sampling directions were all 90. The in-plane resolution and the slice thickness were both 1.25 mm. The diffusion data were reconstructed in the MNI space using q-space diffeomorphic reconstruction (Yeh et al., 2011) to obtain the spin distribution function (Yeh et al., 2010). A diffusion sampling length ratio of 2.5 was used, and the output resolution was 1 mm.

### *Rhesus* macaque *MRI experiments*

A group average template was constructed from a total of 36 animal scans from the PRIMatE Data Exchange (PRIME-DE) (http://fcon_1000.projects.nitrc.org/indi/indiPRIME.html) acquired at 5 different sites including UC Davis, Princeton University, University of Minnesota, Mount Sanai, and American Military University (Milham et al., 2018). Detailed acquisition parameters are described previously. The b-table was checked by an automatic quality control routine to ensure its accuracy (Schilling et al., 2019). The diffusion data were homogenized and reconstructed in the MNI space using q-space diffeomorphic reconstruction (Yeh et al., 2011) to obtain the spin distribution function (Yeh et al., 2010). A diffusion sampling length ratio of 1.25 was used. The reconstructed data were averaged to build a population average diffusion distribution template at 0.5 mm isotropic resolution.

In addition to the population-average template, we also conducted a subject-specific deterministic fiber tractography study in 36 rhesus macaque subjects. The diffusion data were reconstructed using generalized q-sampling imaging (Yeh et al., 2010) with a diffusion sampling length ratio of 1 to obtain the spin distribution function in the native space to enable subject-space fiber tracking.

### Fiber tracking and analysis

We performed diffusion fiber tracking using our proprietary and open-source software DSI Studio (http://dsi-studio.labsolver.org). A deterministic fiber tracking algorithm was implemented using DSI Studio software (Yeh et al., 2013). A seeding region was placed on the whole brain. The anisotropy threshold for the human and rhesus macaque was automatically determined using the default anisotropy threshold in DSI studio. The angular threshold was 75 degrees, and the step size was 1 mm. Tracts with a length shorter than 30 or longer than 300 mm were discarded. The tracts were set to terminate if a total of 10,000 or more tracts were calculated.

The protocol for mapping the anterior commissure for human and rhesus macaque subjects is summarized in Supp. Fig. 1. In the figure, the human HCP 1mm and INDI primate templates were used for visualization purposes. For the human fiber tract protocol, the left and right hemispheres were tracked separately to obtain the orbitofrontal, temporal, and occipital branches of the anterior commissure. The anterior commissure region from the HCP842 tractography atlas (Yeh et al., 2018) was used as a seed region (pink). A small terminating region was placed at the center of the anterior commissure (orange) and a region of interest (ROI) (blue) was placed next to the terminating region on the hemisphere that tracts were obtained. The small terminating region and the large region of interest were created using a sphere from the drawing options in DSI Studio. Supp. Fig. 1a shows how the left orbitofrontal tract (red) was obtained using the left lateral (green) and medial orbitofrontal (orange) lobes. Supp. Fig. 1b shows how the left temporal tract (green) was obtained using the left temporal regions, specifically the left amygdala (blue) and temporal pole (orange). Supp. Fig. 1c shows how the left occipital tract (blue) was obtained using the left_lateral_occipital region (blue). The orbitofrontal and occipital regions both come from the FreeSurferDKT atlas (https://surfer.nmr.mgh.harvard.edu/). The temporal regions come from the Automated Anatomical Labeling (AAL) atlas (Rolls et al., 2020). For all the branches, the right hemispheric fiber tracts were generated similarly except for changing the region of interest and the specified atlas regions to the corresponding right hemisphere. The fiber tracts were cleaned up by removing tracts that did not follow the tract of interest and were apparent to correspond to false tracts.

For the rhesus macaque fiber tract protocol, the left and right hemispheres were tracked separately to obtain the orbitofrontal, temporal, and occipital branches of the anterior commissure. The ROI region (pink) was created using a sphere from the drawing options in DSI Studio to track the anterior commissure in rhesus macaques. It was placed at the center of the sagittal view for obtaining all the fiber tract branches. Supp. Fig 1d, 1e, and 1f show the representative fiber tracts for the orbitofrontal, temporal, and occipital branches of the rhesus macaque anterior commissure, respectively. The branches were cleaned up by removing tracts that did not follow the tract of interest and were apparent to correspond to false tracts.

We further calculated the volume fraction of the anterior commissure component and the corpus callosum component of the redundancy circuits. The volume fraction was calculated by dividing the volume of the tracks by the brain volume. We removed three outliers from the human data and six from the rhesus macaque whose values were more than three scaled median absolute deviations for both the anterior commissure and corpus callosum tracts. A two-sample unequal variances t-test was used to test the differences in the volume fractions of the redundancy circuits of the commissural pathways of the human and rhesus macaque brains. We also calculated the component percentage of the anterior commissure and corpus callosum component in each redundancy circuit and tested the differences using a two-sample unequal variances t-test.

### Connectogram

For the human subjects, the connectivity matrix function in DSI studio was used to create matrices representing the number of fibers terminating within regions of the FreeSurferDKT atlas in reference to the terminating region used for fiber tracking. For the rhesus macaque subjects, the CIVM rhesus macaque atlas (Calabrese et al., 2015) was used in reference to the ROI used for fiber tracking of the rhesus macaque anterior commissure. In creating the connectivity matrix for both the human and rhesus macaque subjects, the ROIs were specified as pass regions. Two connectivity matrices were created per subject corresponding to the left and right hemispheres. In total, 60 connectivity matrices were generated for the human subjects and 72 connectivity matrices were generated for the rhesus macaque subjects. The connectivity values were normalized and averaged across all the human and rhesus macaque subjects, respectively, to generate the connectogram.

### Cadaveric dissections

The Klingler method (Ludwig and Klingler, 1956) was used for dissection of the human and rhesus macaque anterior commissure. Two human (4 hemispheres) and two rhesus macaque (4 hemispheres) brains were prepared for dissection by first fixing them in a 10% formalin solution for 2-4 weeks. After fixation, the brains were frozen at −16C for 2-3 weeks, and dissection commenced 24h after the specimens were thawed. The first steps consisted of careful removal of the arachnoid and superficial vessels followed by a step-wise, superficial-to-deep dissection. Wooden spatulas were used to remove successive layers of gray matter followed by white matter to reveal the anterior commissure. In between dissections, specimens were stored in 10% formalin solution. High-quality photographs were taken before dissection to identify sulci and gyri and then as the dissection proceeded to localize other white fiber pathways. Photographs were inverted using Pixlr Editor for another method of visualizing the fiber pathways. The inverted images for the human and rhesus macaque dissections are included in the supplementary material (Fig. S4 and S5, respectively) (Sevandersson, 2020).

