## Supplementary material for "Redundancy Circuits of the Commissural Pathways in Human and Rhesus Macaque Brains"

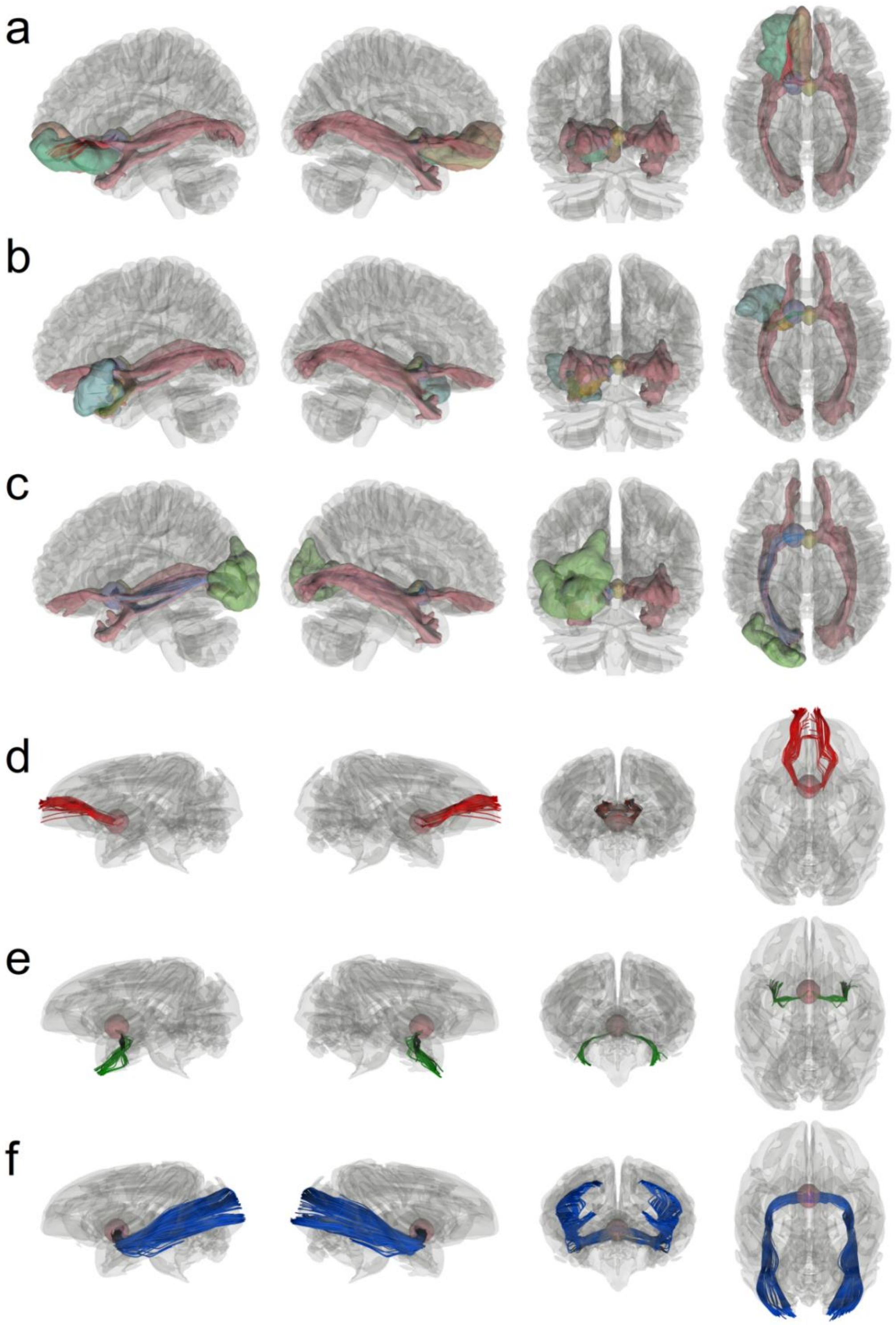

**Supplementary Figure 1:** Protocol for mapping the anterior commissure for human and rhesus macaque subjects

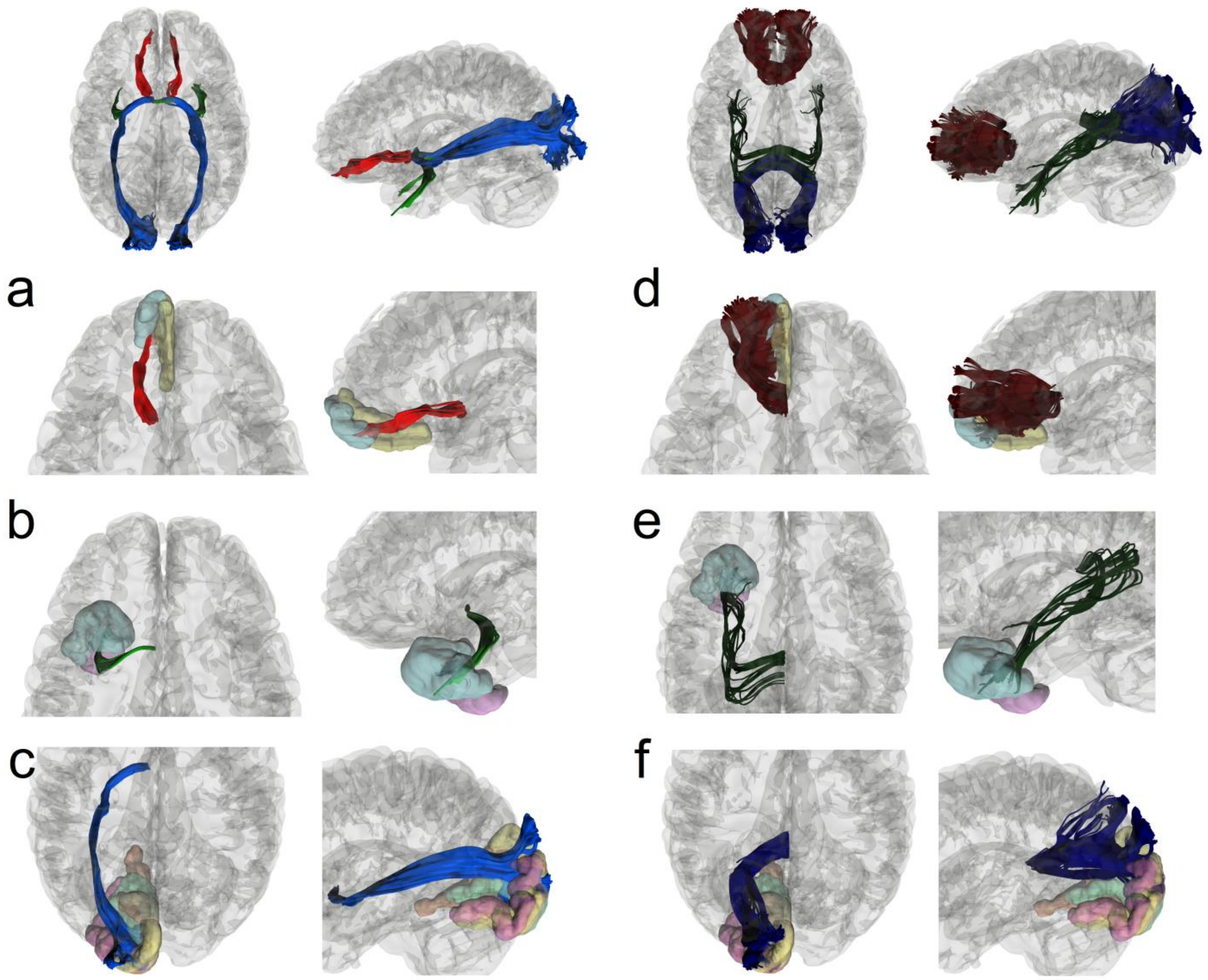

**Supplementary Figure 2:** Anterior commissure and corpus callosum visualization in the human brain

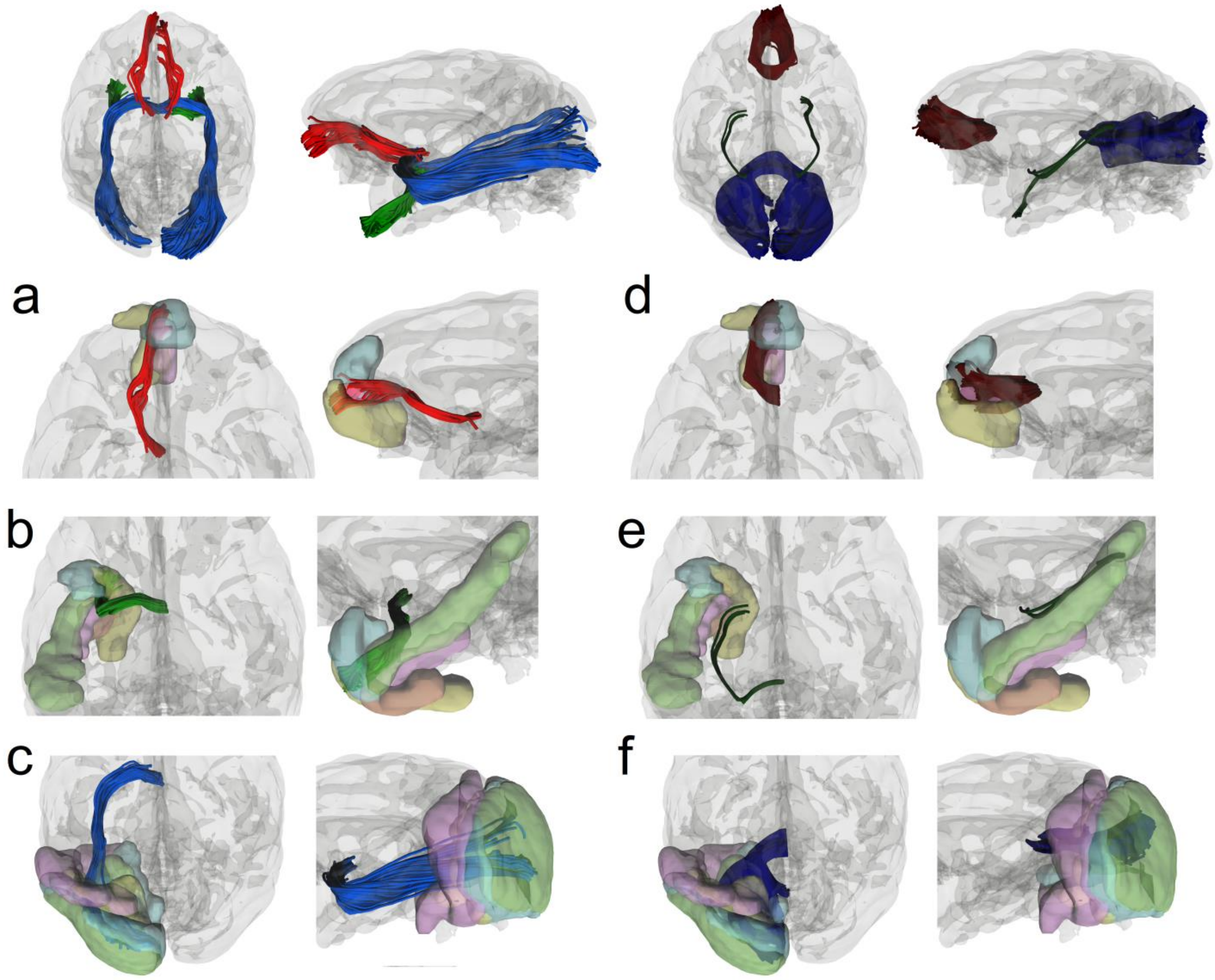

**Supplementary Figure 3:** Anterior commissure and corpus callosum visualization in the rhesus macaque brain

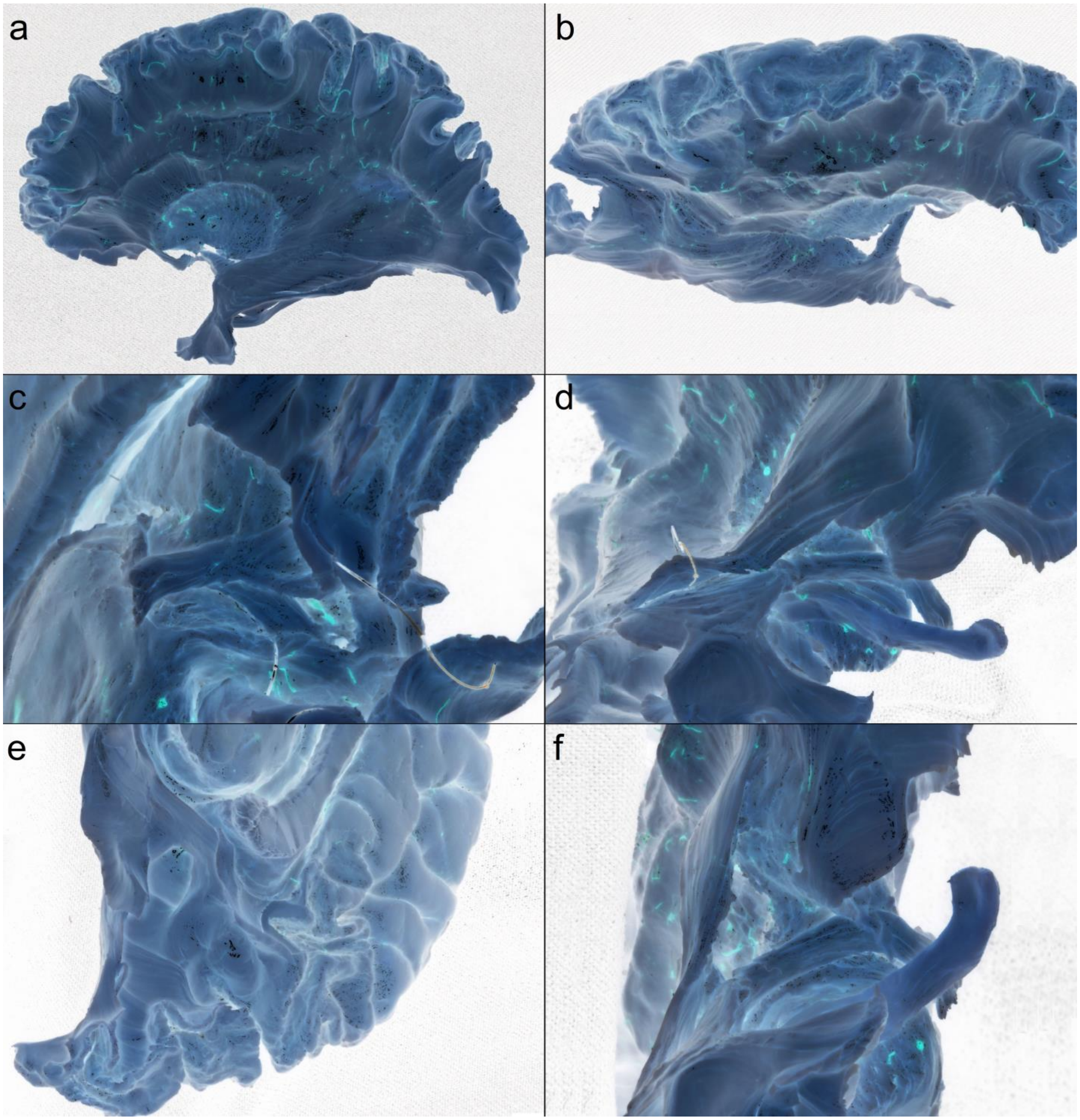

**Supplementary Figure 4:** Inverted human cadaveric dissection images

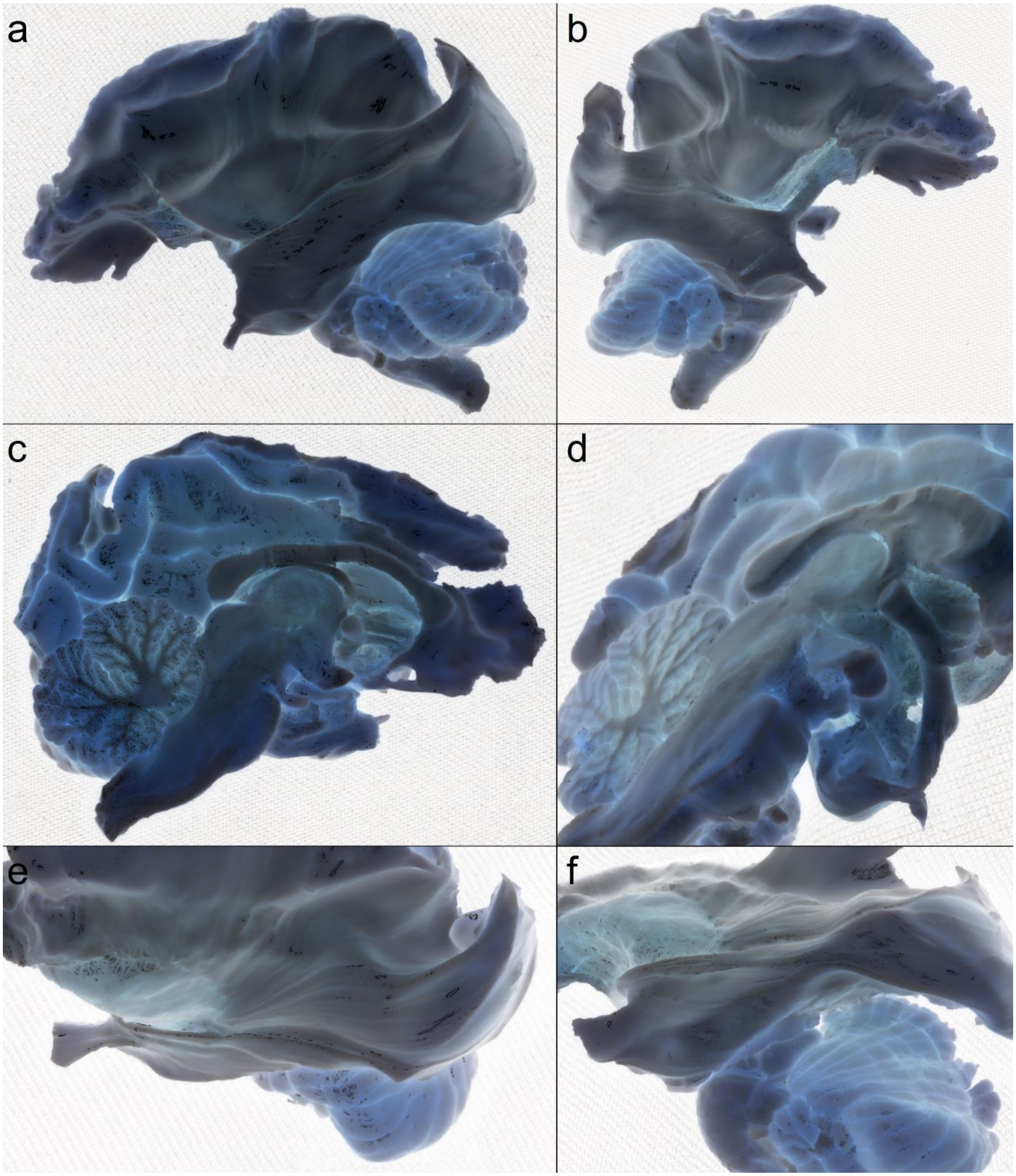

**Supplementary Figure 5:** Inverted rhesus macaque cadaveric dissection images

**Supplementary Table 1**: Regions the anterior commissure projects to in the human and rhesus macaque brain

| | Occipital | Orbitofrontal | Temporal |
|---|---|---|---|
| Human | V1<br>V2<br>V3<br>V4 | 10v<br>10pp | TGd<br>TGv |
| Rhesus macaque | V1<br>V2<br>V3<br>V4 | 10M<br>10V<br>14M | IPa<br>TLR (R36)<br>TE1<br>TPO<br>TPPro |

**V1:** primary visual cortex

**V2:** secondary visual cortex

**V3:** tertiary visual cortex

**V4:** quaternary visual cortex

**10v:** area 10 ventral

**10pp:** polar 10p

**TGd:** area TG dorsal

**TGv:** area TG ventral

**10M:** area 10 of cortex medial part

**10V:** area 10 of cortex ventral part

**14M:** area 14 of cortex medial part

**IPa:** intraparietal sulcus associated area in the superior temporal sulcus

**TLR(R36):** area TL rostral part (area 36R)

**TE1:** temporal area

**TPO:** temporal parietoccipital associated area in sts

**TPPro:** temporopolar prisocortex

**Supplementary Table 2:** Abbreviations used in the dissection images

| Abbreviations | Brain Regions |
|---|---|
| AC | Anterior commissure |
| AnsPed | Ansa Peduncularis |
| VA | Ventral anterior nucleus |
| Calc Fiss | Calcarine Fissure |
| CN | Caudate Nucleus |
| CC | Corpus callosum |
| CB | Cerebellum |
| CR | Corona Radiata |
| FB | orbitofrontal branch |
| F | Fornix |
| HP | Hippocampus |
| IFOF | Inferior Front-Occipital Fasciculus |
| Lent | Lentiform nucleus |
| M | Mammillary bodies |
| OB | Occipital branch |
| Opt | Optic nerve |
| POSulc | Parietooccipital Sulcus |
| P | Pons |
| PreCun | Precuneus |
| PT | Putamen |
| RectG | Rectus Gyrus |
| SLF | superior longitudinal fasciculus |
| TH | Thalamus |
| TB | Temporal branch |
| UN | Uncinate fasciculus |
| V1 | primary visual area |
| V2 | secondary visual area |
| V3 | tertiary visual area |
